# Impaired HIV-specific T-cell responses in HIV and *S. mansoni* coinfected Ugandans

**DOI:** 10.1101/2022.11.01.514652

**Authors:** Andrew Ekii Obuku, Ashish Sharma, Anjellina Rukundo, Matthew Odong, Betty Auma, Jennifer Serwanga, Ronald Nkangi, Joseph Okello, Zahara Nakitto, Moses Joloba, Giuseppe Pantaleo, Pontiano Kaleebu, Rafik Sekaly

## Abstract

Fishing communities surrounding Lake Victoria in Uganda show an HIV-1 prevalence of 28% and incidence rates of 5%. More than 50% of the fishermen on the shores of Lake Victoria are infected with *S. mansoni*. Fishermen are more likely to die of AIDS related illness than farmers in the Lake Victoria region.

Using polychromatic flow cytometry and mesoscale discovery platform, HIV specific and non-specific responses were measured and compared within individuals when HIV and *S. mansoni* coinfected and after the *S. mansoni* was cleared.

Sixty-two unique clusters of cells in the UMAP space were identified after stimulations with GAG PTE POOL-1 and GAG PTE POOL-2 independently. However, the frequency of only three clusters is significantly higher after S. *mansoni* clearance. In addition, *S. mansoni* infection is associated with higher IL-9 and IL-10 and lower IL-15 in HIV and *S. mansoni* coinfected individuals. IL-9 concentration at enrolment visit predicts CD4 decline.

*S. mansoni* infection negatively affects HIV specific and non-specific immune responses in HIV and *S. mansoni* coinfected individuals.

## Introduction

Although mixed infections occur more often than not, there is limited evidence for the beneficial effects of their interactions. Interactions between the coinfecting pathogens and the host may alter the pathogenesis and transmission of one or both pathogens [1], disease progression [2] and control of multiple diseases. Immunological effects of mixed infection include by stander immune activation in the absence of cognate antigens [3] and cellular attrition [4].

Globally, fishing communities have enormous challenges of access to health services [5]. The outcome of which, is the high HIV prevalence and incidence rates in Uganda [6, 7]. They also have high *Schistosoma mansoni* prevalence rates [8]. With significant geographical overlap, coinfection of HIV and *S. mansoni* is very common in the fishing communities.

Although the number of deaths as a result of schistosomiasis alone is low, there is a great deal of unacknowledged morbidity [9]. Among the possible subtle effects of schistosomiasis, an unresolved question is the extent to which schistosomiasis influences the host response to vaccinations and other infections such as HIV and how this affects the rate of progression to AIDS.

Few studies have directly investigated the association between *S. mansoni* infection and HIV progression to AIDS. However, multiple studies have shown that the plasma HIV RNA level is predictive of both HIV disease progression [10, 11] and risk of transmission of HIV to sexual partners [12] and that the treatment of *S. mansoni* infection with praziquantel (PZQ) reduced the rate of CD4 T cell decline in previously HIV and *S. mansoni* coinfected participants [13]. While treatment for schistosomiasis is clearly not able to substitute for antiretroviral therapy, it may possibly be able to slow HIV disease progression.

Cellular immune responses including CD8 T cells, CD4 T cells, B cells and NK cells are thought to play a significant role in the control of HIV infection [14, 15], and coinfection with *S. mansoni* may affect the quantity and quality of the cellular anti HIV specific immune responses [16], since as a single infection, adult schistosomes suppress immunity by modulating immune system components, including T helper (Th1) and interferon signaling [16]. Scanty data is available to understand whether this immune suppression occurs in a coinfection of *S. mansoni* and HIV infection.

Individually, HIV infection induces a predominant Th1 biased immune response and *S. mansoni* induces a predominant Th2 biased immune response. If Th1 and Th2 are mutually exclusive, the predominant Th phenotype in individuals coinfected with HIV and *S. mansoni* has not been well characterized. In addition, the effect of *S. mansoni* infection on HIV-specific immune responses in HIV and *S. mansoni* coinfected individuals have also not been well characterized.

Indeed, in a mixed infection of HIV and *S. mansoni* this study, a) the frequency of HIV specific T cells and IL-10 responses were significantly lower, however, IL-9 and IL-15 responses were significantly higher, during the mixed infection : b) Increase in the egg counts increased with increase in IL-10 concentration: c) IL-9 concentration at enrolment predicts a significant loss of CD4 count at exit visit: d) IL-9 concentration at exit visit predicts increase in plasma viral load at exit visit.

## Methods

### Ethics statement

The Institutional Ethics Review Board of the Uganda Virus Research Institute (GC/127/12/02/01) and the Uganda National Council of Science and Technology (HS1141) approved the study. All study participants provided written informed consent.

### Study group

Antiretroviral treatment (ART)-naïve HIV and *S. mansoni* co-infected participants without hookworm, *Ascaris lumbricoides, Strongyloides stercoralis, Giardia lamblia, Trichuris trichiura, Clonorchis sinenis, Hymenolepis diminuta* or *Fasciolopsis buski* were treated with a single praziquantel dose (40mg/kg) once every three months for 60 weeks (exit visit), the first two doses were administered one week apart. Participants who had cleared *S. mansoni* infection (i.e. negative by both circulating anodic antigen (CAA) and Kato-Katz (K.K) tests) by week 48 were selected for the immunology study. Samples collected at the enrolment visit when participants were HIV and *S. mansoni* coinfected and week 60 (exit visit) when participants were HIV infected and *S. mansoni* negative (i.e., negative by both CAA and K.K tests) were selected.

### Flow cytometry staining assay

Cryo-preserved peripheral blood mononuclear cells (1×10^6^) were thawed, rested for 6 hours and stimulated overnight in 1 ml of complete medium (RPMI 1640 with Glutamax, Gibco) containing 10% fetal bovine serum (FBS; PAA, Austria), 100μg/ml penicillin, 100 units /ml streptomycin (Lonza, Switzerland) in the presence of Golgiplug (1μl/ml, BD) and 1 μg/ml of potential T cell epitopes (PTE) peptide pools (GAG PTE POOL-1 and GAG PTE POOL-2)[17]. Staphylococcus enterotoxin B (SEB; Sigma) stimulation (200ng/ml) was used as a positive control. Cells with medium alone was used as a negative control. At the end of the stimulation period, cells were stained for 20 minutes in the dark at room temperature using Aqua LIVE/DEAD cell stain (Invitrogen), washed and stained for an additional 30 minutes with anti-human CD3 APC-H7, CD4 PE-CF594, CD8 Pacific Blue (all from BD, Biosciences) at 4°C. The cells were then fixed and permeabilized (Cytofix/Cytoperm, BD) at RT for 30 minutes and then stained at with anti-human TNF-α PE-Cy7, IL-2 PerCP Cy5.5, IFN-γ APC (all from BD, Biosciences), anti-human TLR-7 PE (from R & D systems), and anti-human Perforin FITC (from GenProbe). The stained cells were washed twice with buffer containing 98% PBS and 2% FBS. Cellfix (BD, San Diego, CA, USA) was added and 0.6-1×10^6^ lymphocyte-gated events were immediately acquired on an LSRII SORP (BD, San Diego, CA, USA).

### Cytokine quantification using mesoscale discovery platform (MSD)

The assay was performed following the manufacturer’s instructions. Briefly, each biotinylated antibody was added to an assigned linker, vortexed and incubated at room temperature for 30 minutes. Stop solution was added, vortexed and further incubated for 30 minutes at room temperature. IFN-γ, IL-1β, IL-2, IL-4, IL-6, IL-8, IL-10, IL-17A, TNF-α, IL-9 (plate 1), IFN-α2a, IL-7, IL-15, IL-18, IL-21, IL-22, IL-27, IL-29/IFN-1, IL-33, IFN-β, (plate 2), TGF-β 1, TGF-β 2 and TGF-β 3 (plate 3) antibody linker pairs were combined and coated overnight on three separate plates on a shaker at 2-8°C (Meso Scale Diagnostics, LLC, Rockville, Maryland, USA).

Plasma samples were thawed in a water bath maintained at 37°C and aliquoted into plates in duplicate wells. For TGF-β detection, the plasma samples were treated with 1M hydrochloric acid and neutralized by NaOH/HEPES solution.

The plasma samples and standards were added to the plates and incubated for 1 hour at room temperature on a plate shaker at 750 rpm. Detection antibody was added after 3 washes using PBS-T (1x PBS + 0.05% Tween 20). Read buffer T was added and plates were read using mesoscale discovery reader (Meso Scale Diagnostics, LLC, Rockville, Maryland, USA).

### UMAP, PhenoGraph and K means analysis

FCS3.1 data files were imported into Flowjo software version 10.7 (Flowjo LLC). All samples were compensated electronically, gated (supplementary figure 1) and exported. To exclude doublets, data on FSC-A and FSC-H were used. Aqua positive cells (dead cells) were excluded. The unstimulated control for each respective cytokine was used to set the gates for the GAG PTE POOL-1 and GAG PTE POOL-2 responses at twice or more the response of the unstimulated control). Individual cytokine/perforin or TLR7 expressing cells were Boolean gated using the OR function.

**Figure 1.**
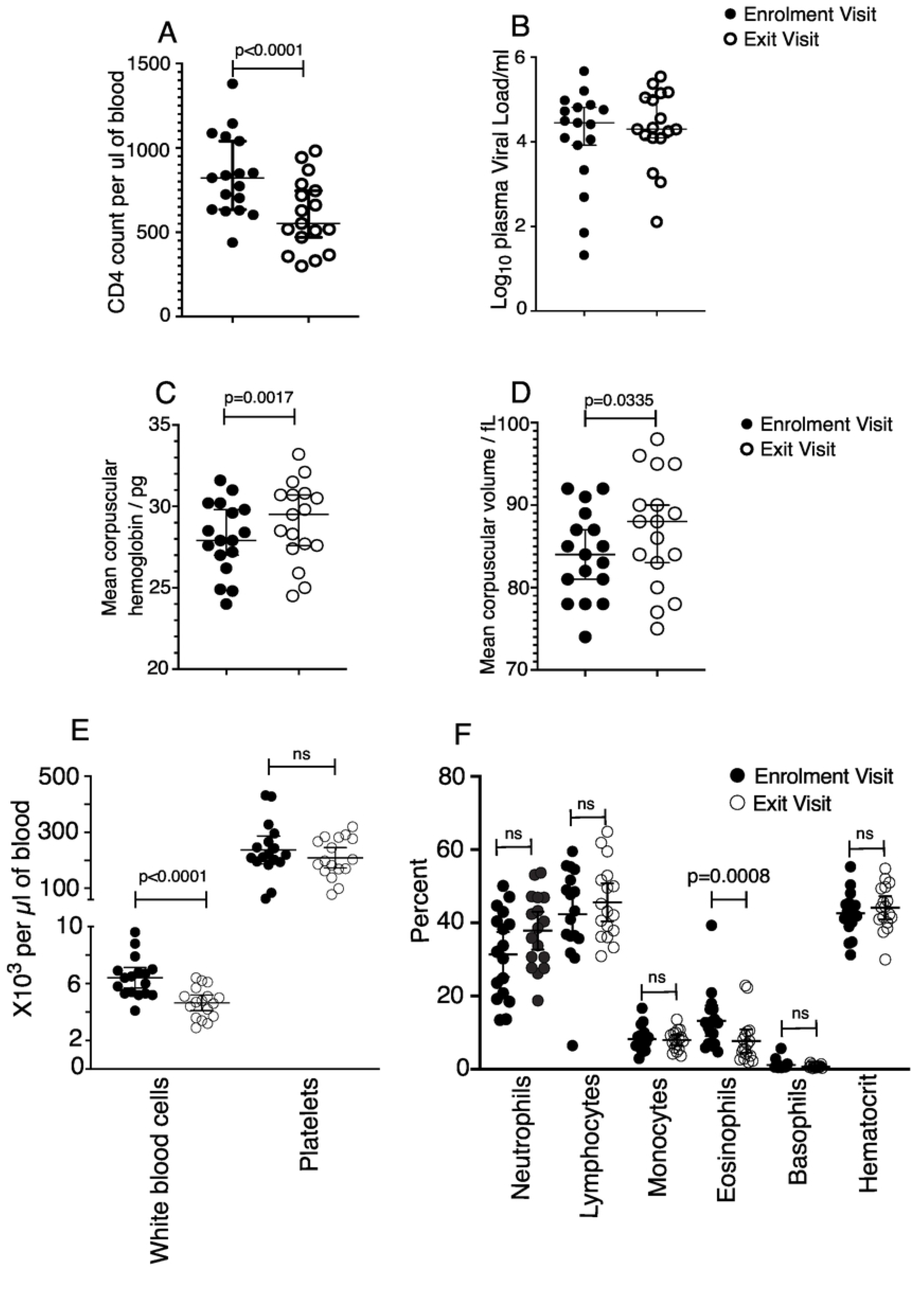
CD4, plasma Viral Load and complete blood count at enrolment and exit visits. Each dot represents results for a participant. The horizontal lines represent the mean and 95% confidence interval (C.I). Paired student t test (A), Wilcoxon matched pairs signed rank test (B) and RM one way ANOVA with Sidak’s multiple comparison test (C, D, E, F) were used to compare the mean between enrolment and exit visits. ns represents nonsignificant. n=17.

umap-learn version 0.4.6 [18] was used in the dimensionality reduction of equal number of cells from each CSV file (8,757 from each of GAG PTE POOL-1 file and 18,383 from each of GAG PTE POOL-2 file selected randomly). The file with the lowest number of events determined the number of events contributed by each file for a particular stimulation. Clusters of phenotypically similar cells were detected using PhenoGraph [19] in Rphenograph version 0.99.1. PhenoGraph clustering was performed using UMAP1 and UMAP 2 coordinates. IFN-γ, IL-2, TNF-α, perforin, CD3, CD4, CD8 and TLR7 intensities were used in dimensionality reduction and clustering.

To validate Louvain community detection clustering, the median fluorescence intensity of each marker (IFN-γ, IL-2, TNF-α, perforin, CD3, CD4, CD8 and TLR7) were also clustered using K means.

The frequency of cells in each cluster identified using Louvain community detection was compared between enrolment and exit visits.

### Statistical analysis

Differences between matched groups were compared using paired T test or Wilcoxon matched-pairs signed rank test. Nonparametric tests were used if the data were not normally distributed. Shapiro-Wilk normality test was used to test if the data is normally distributed.

All analyses were performed using R studio or GraphPad prism version 8.4.3.

## Results

### Bio-medical characteristics of study participants

Seventeen participants who were coinfected with HIV and *S. mansoni* (HIV+SM+) at enrolment into the study and had cleared *S. mansoni* infection (HIV+SM-) at exit visit were selected. Four participants were females. The mean age of the participants at enrolment was 29 years (range 18-41 years). Although plasma viral load (pVL) was not significantly different between enrolment and exit visits (Median pVL at enrolment and exit visits were 28,084 HIV RNA copies/ml and 19,834 HIV RNA copies/ml, respectively p= 0.7819: Wilcoxon matched pairs signed rank test) **figure 1B**, the CD4 count was significantly lower by 232.7 cells per μl of blood at the exit visit (p=<0.0001, paired t test: 95% C.I. -319.1 to -146.3) **figure 1A**. The number of white blood cells per μl of blood (6.406 X10^3^ per microliter of blood at enrolment and 4.653 X10^3^ per microliter of blood at exit visit) and percent of eosinophils in blood (13.28% at enrolment and 7.753% at exit visit mean difference 5.529 95% CI of difference 2.217-8.842) were also significantly lower at the exit visit (adjusted p value <0.0001 and p=0.0008, respectively, RM one way ANOVA with Sidak’s multiple comparison test) **figure 1E and 1F**. Although within the normal ranges (76.0-96.0fL for mean corpuscular volume (MCV) and 27.0-32.0pg for mean corpuscular hemoglobin (MCH)), the MCV (83.94fL and 86.82fL at enrolment and exit visits, respectively) and MCH (28.05 pg and 29.04 pg at enrolment and exit visits, respectively) were significantly higher at the exit visit (adjusted p value p=0.0335 and p=0.0017, respectively, RM one way ANOVA with Sidak’s multiple comparison test) **figure 1D and 1C**.

### Frequency of unique HIV specific T cell subsets were enhanced on *S. mansoni* clearance

To identify HIV specific cells that are impacted by *S. mansoni* coinfection, PBMC were stimulated with GAG PTE POOL-1 and GAG PTE POOL-2 peptides. In response to GAG PTE POOL-1 stimulation, PhenoGraph clustering revealed 30 unique clusters of cells in the UMAP space **figure 2A and 2B**. However, only the frequency of cells in clusters 11 (Median =0.006737% and 0.02084% at enrolment and exit visits, respectively) and 16 (Median =0.009992% and 0.02341% at enrolment and exit visits, respectively) were significantly higher at the exit visit compared to the enrolment visit (p=0.0023 and p=0.0134: Wilcoxon matched pairs signed rank test) **figure 3A and 3B**.

**Figure 2.**
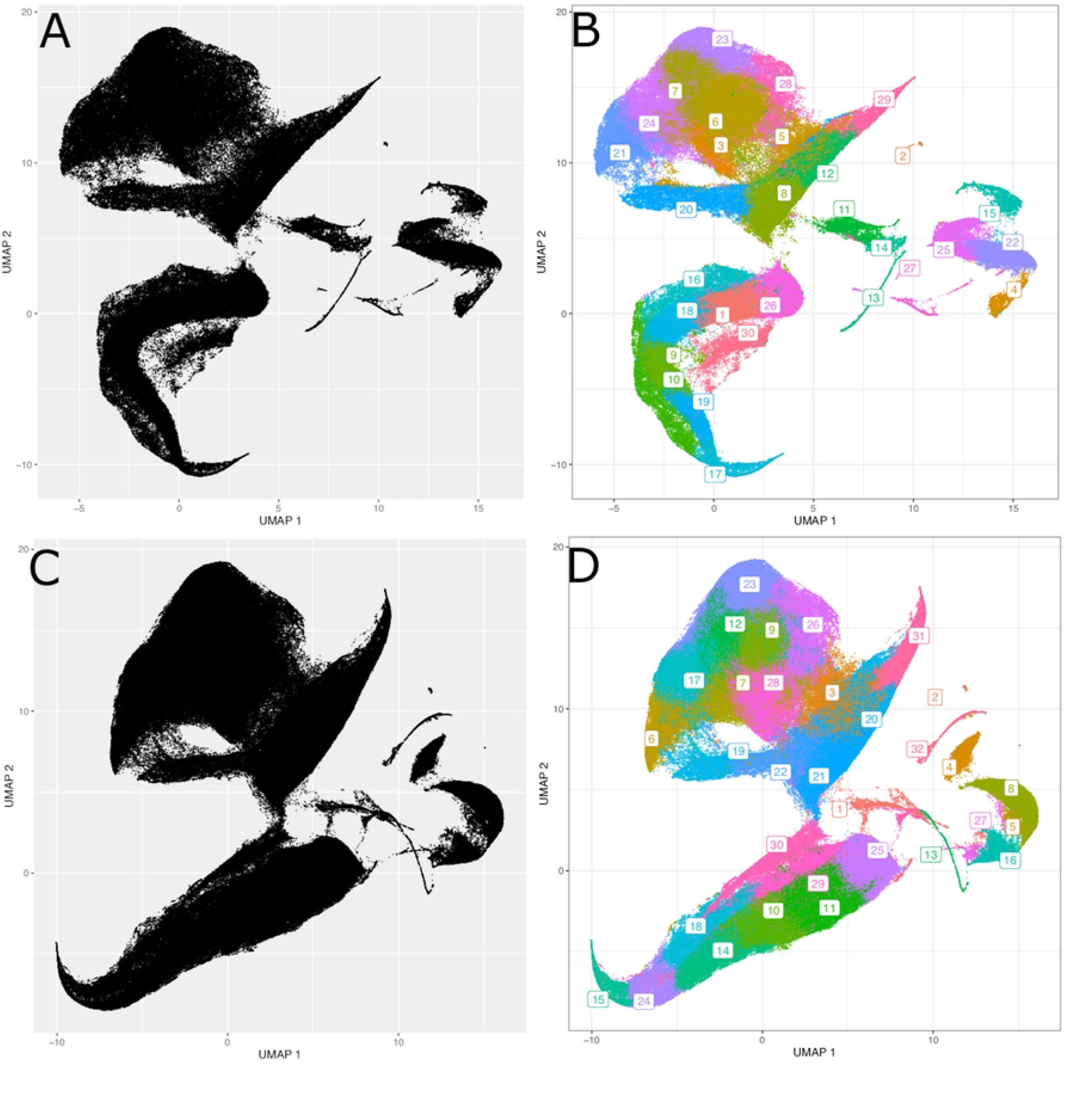
Clustering using Louvain community detection followed by projection on the UMAP map with T cells from enrolment and exit visits. Each dot represents a cell from either enrolment or exit visit. Cells were stimulated with Gag PTE POOL-1 (A and B) or Gag PTE POOL-2 (C and D). Figure A and C shows UMAP scatter plots. Figures B and D shows clusters of phenotypically similar cells detected using Louvain community detection. The numbers 1 to 32 are the different cell clusters identified within each dataset. Cells with identical colors belong to the same cluster.

**Figure 3.**
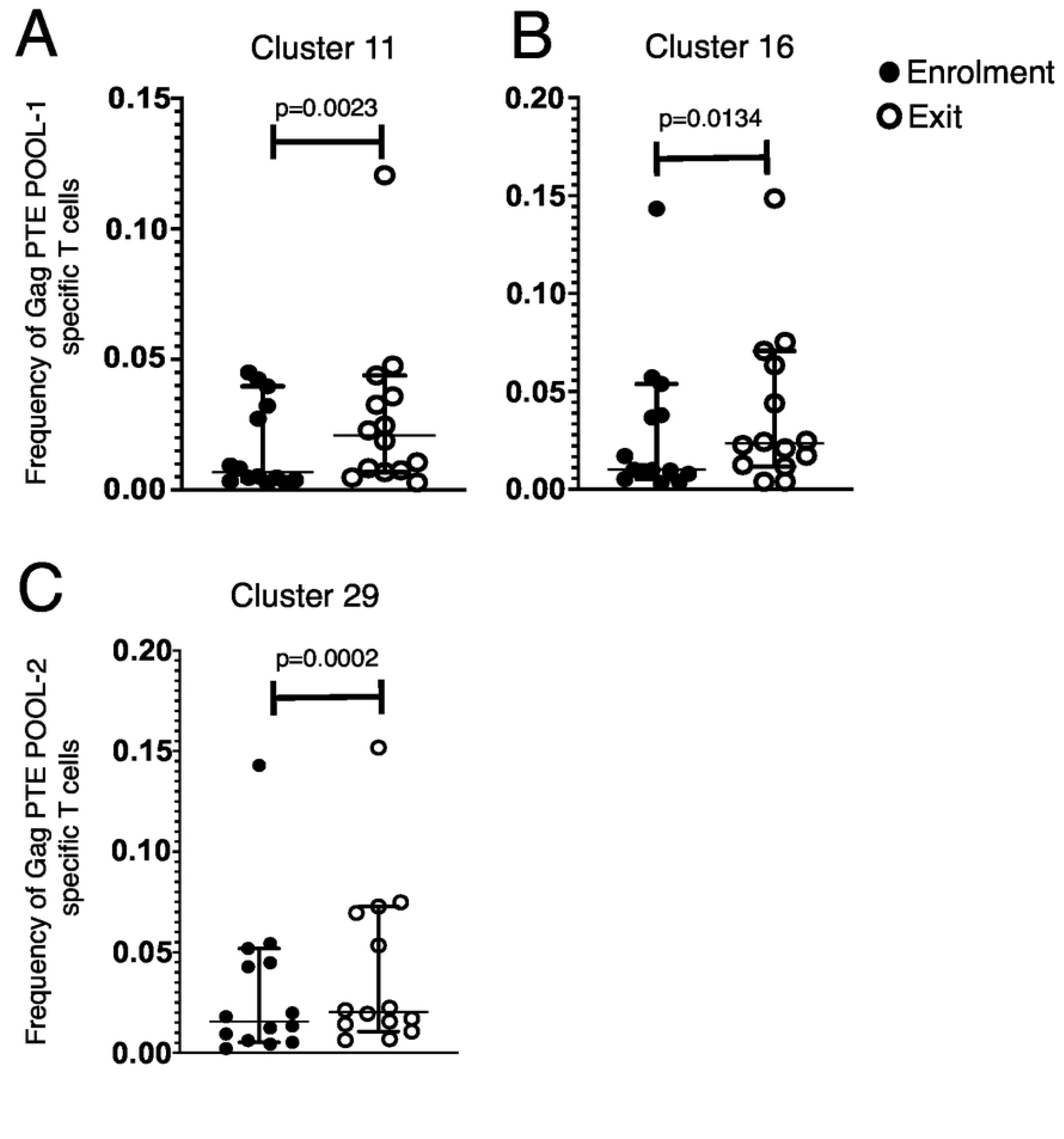
Frequency of Gag PTE POOL-1 and Gag PTE POOL-2 specific T cells between enrolment and exit visits from clusters 11, 16 and 29. Cells in clusters 11 and 16 are Gag PTE POOL-1 stimulated whereas cells in cluster 29 are Gag PTE POOL-2 stimulated. The horizontal lines represent the median and 95% confidence interval (C.I) Wilcoxon matched pairs signed rank test was used to compare the frequency of Gag specific T cells between enrolment and exit visits. n=14

Cells in cluster 11 had high expression intensities of IFN-γ intermediate expression intensities of TNF-α, CD8 and CD3 and low expression intensities of perforin, TLR7, CD4, and IL-2 (figure **4A-I**). Cells in cluster 16 had intermediate expression intensities of CD8 and perforin and low expression intensities of IFN-γ, TNF-α, CD3, TLR-7, CD4 and IL-2 (**figure 4A-I**).

**Figure 4.**
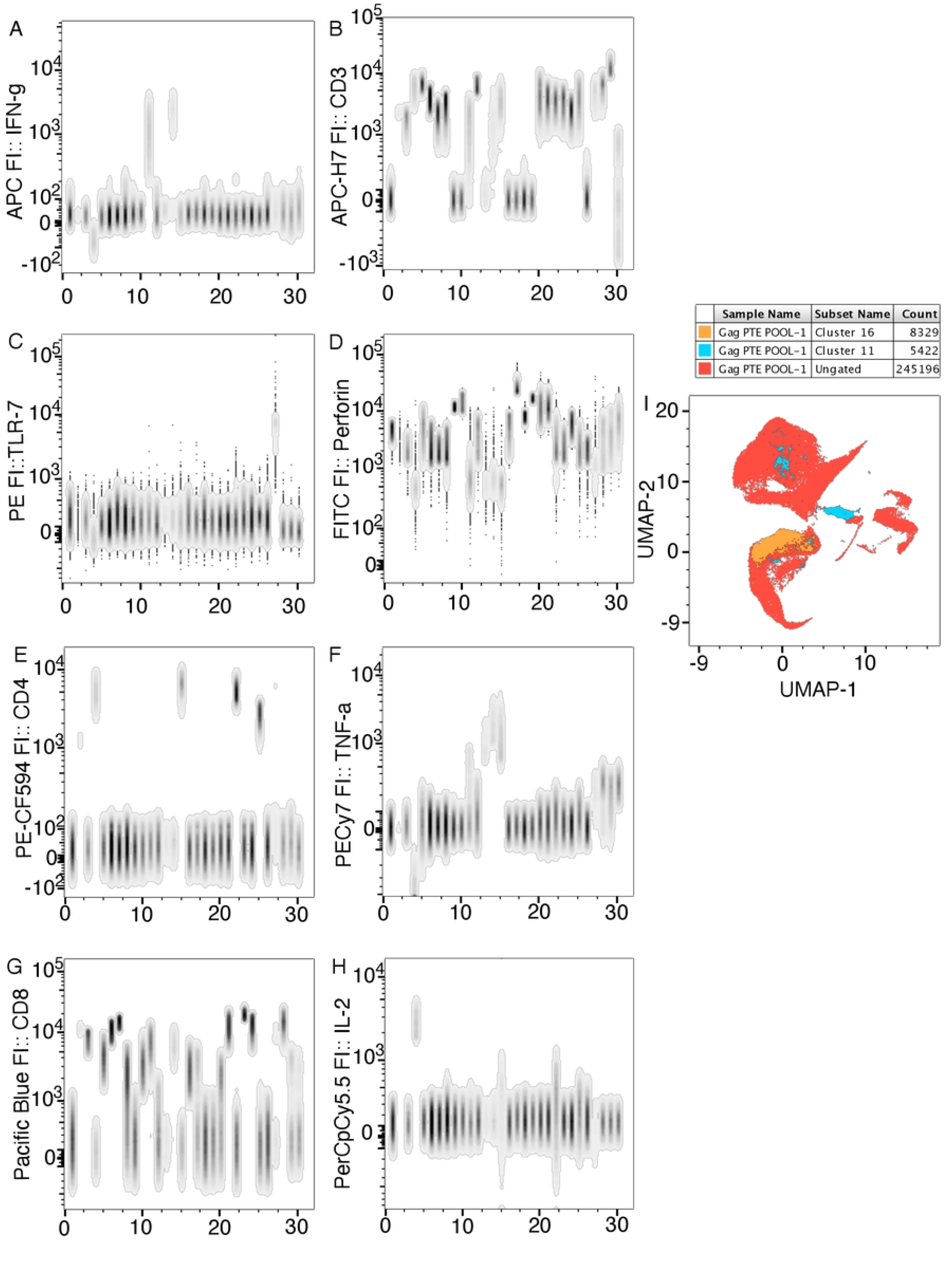
Expression intensity of markers for cells in cluster 11 juxtaposed on all the cells in the experiment.

In response to GAG PTE POOL-2 stimulation, PhenoGraph analysis revealed 32 unique clusters of cells in the UMAP space (**figure 2C and 2D)**. However, only the frequency of cells in cluster 29 (Median =0.01569% and 0.02037% at enrolment and exit visits, respectively) were significantly higher at the exit visit compared to the enrolment visit (p=0.0002: Wilcoxon matched pairs signed rank test figure **3C**).

Cells in cluster 29 had intermediate expression intensities of CD8 and perforin and low expression intensities of IFN-γ, TNF-α, CD3, TLR7, CD4 and IL-2 (**figure 5A-I**).

**Figure 5.**
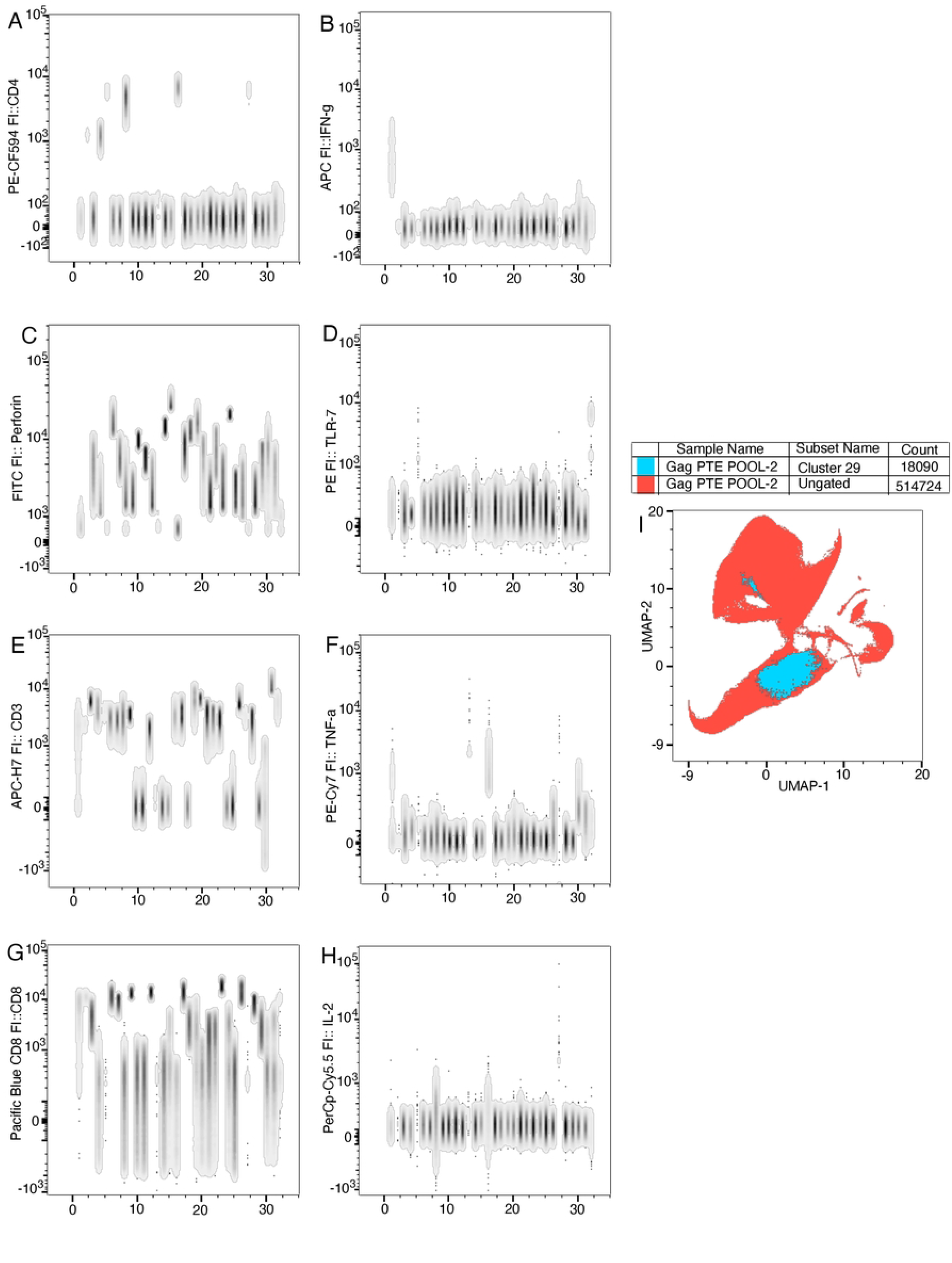
Expression intensity of markers for cells in cluster 16 juxtaposed on all the cells in the experiment.

Two mega clusters of cells were identified in response to GAG PTE POOL-1 stimulation using K means clustering (**figures 6A and 7A**). Mega cluster 1 contained clusters 30, 10, 16, 17, 19, 26, 1, 9, and 18 whose cells preferentially express high levels of perforin, intermediate levels of TLR7, IFN-γ, TNF-α, CD4, IL-2 and low levels of CD3 and CD8. Mega cluster 2 contained clusters 15, 22, 25, 11, 14, 13, 27, 12, 29, 20, 5, 8, 2, 21, 24, 28, 3, 6, 7 and 23. The mega cluster 2 contains four sub mega clusters: sub mega cluster A made up of clusters 15, 22, 25 whose cells express high levels of CD4, intermediate levels of IL-2, CD8, TLR-7, IFN-γ, TNF-α and low levels of CD8 and perforin: sub mega cluster B made up of clusters 11, 14, 13, 27 whose cells preferentially express high levels of IFN-γ and TNF-α and low levels of perforin: sub mega cluster C made up of clusters 12, 29, 20, 5 and 8 whose cells preferentially express intermediate levels of IL-2, CD8, TLR-7, IFN-γ, TNF-α, CD8, perforin and CD4: sub mega cluster D made up of clusters 2, 21, 24, 28, 3, 6, 7, 23 whose cells preferentially express high levels of CD3 and CD8, low levels of perforin, CD4 and IL-2 and intermediate levels of TLR7, IFN-γ and TNF-α.

**Figure 6.**
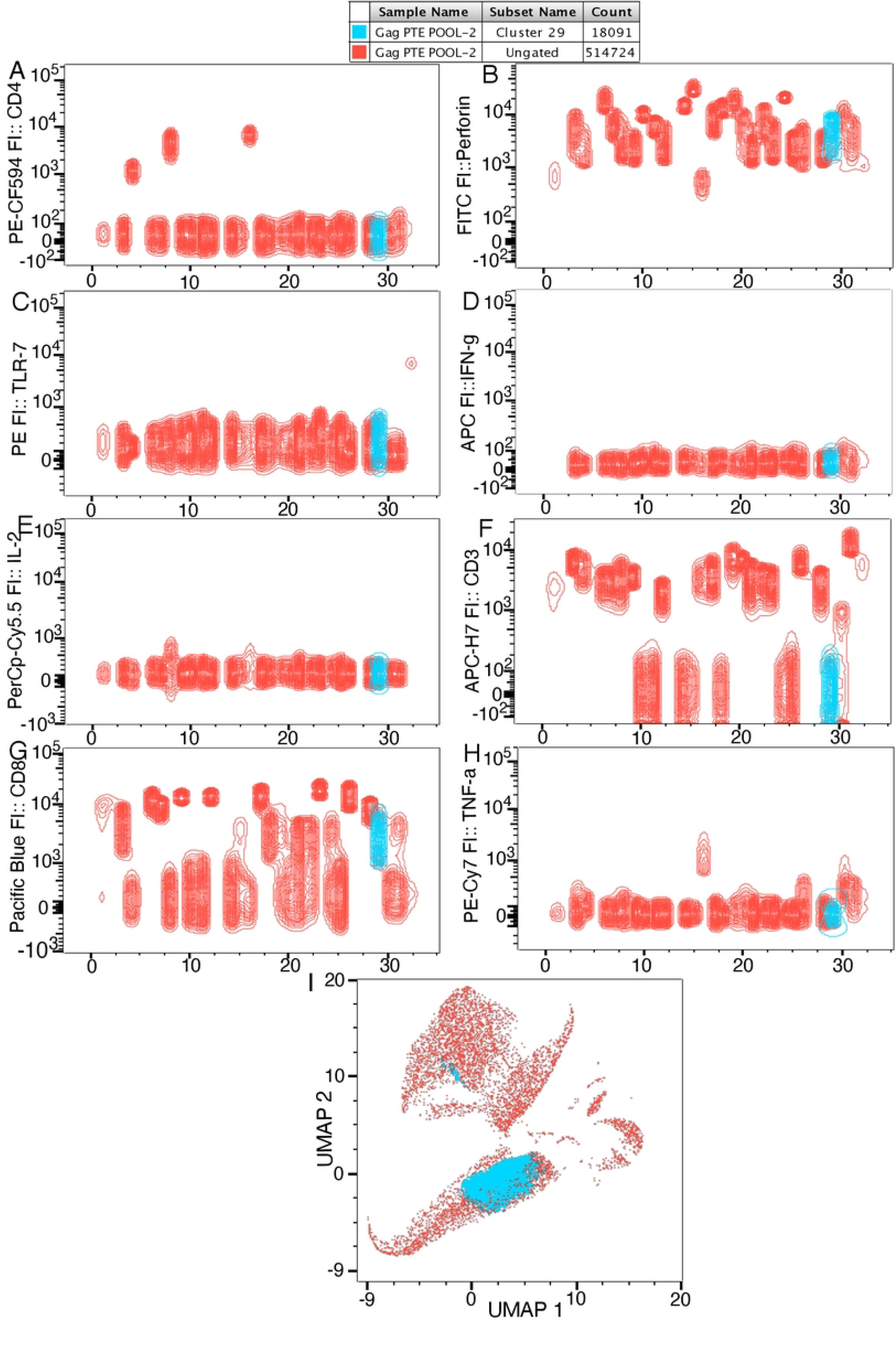
Expression intensity of markers for cells in cluster 29 juxtaposed on all the cells in the experiment.

**Figure 7.**
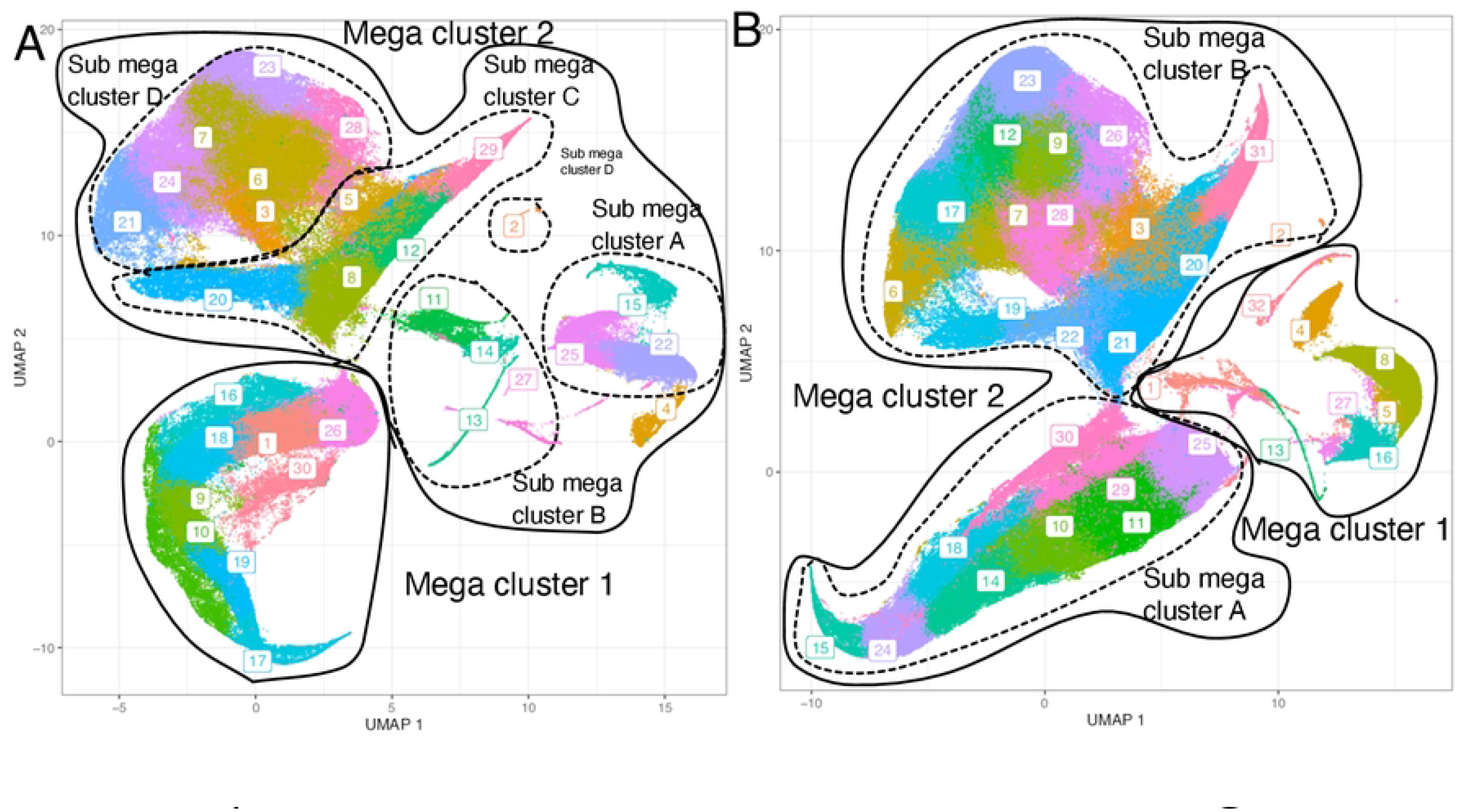
Marker expression levels for each cell cluster after Gag PTE POOL-1 (A) or Gag PTE POOL-2 (B) stimulation. K means clustering was done on the mean fluorescence intensity of each cell to show the relationships between the clusters. The numbers at the bottom of each plot represents the cluster number identified by Louvain community detection algorithm. The key on the right shows the expression levels of the respective marker after Gag PTE POOL-1 or Gag PTE POOL-2 stimulation.

In response to GAG PTE POOL-2 stimulation, two mega clusters 1 and 2 were identified (**figures 6B and 7B**). Mega cluster 1 consists of clusters 27, 13, 5, 16, 4, 8, 1, 32. The cells in mega cluster 1 predominantly expressed high levels of TNF-α and CD4, intermediate levels of IL-2 and CD3, low levels of IFN-γ, TLR-7, CD8 and perforin. Mega cluster 2 consists of two sub mega clusters A and B. Cells in mega sub cluster A predominantly expressed intermediate levels of perforin, low levels of CD3, CD8, TNF-α, CD4, IL-2, IFN-γ and TLR-7. Cells in mega sub cluster B predominantly expressed intermediate levels of CD3 and CD8 and low levels of perforin, TNF-α, CD4, IL-2, IFN-γ and TLR-7.

### *S. mansoni* infection was associated with higher IL-9 and IL-10 and lower IL-15 in HIV and *S. mansoni* coinfected individuals

Cytokine concentrations in plasma were compared at enrolment and exit visits. The IL-9 (median =0.5692 pg/ml and 0.2117 pg/ml) and IL-10 (median =0.6063 pg/ml and 0.4209 pg/ml) concentrations were significantly lower at the exit compared to the enrolment visit, respectively (p=0.0001 and p=0.0067, respectively: Wilcoxon matched pairs signed rank test). However, IL-15 concentration was significantly higher at the exit visit compared to the enrolment visit (median =1.720 pg/ml and 1.909 pg/ml at enrolment and exit visits, respectively, p=0.0034: Wilcoxon matched pairs signed rank test) **Figure 8A, B, C**. To understand the relationship between *S. mansoni* burden and IL-9, IL-10 and IL-15 concentrations, linear regression analyses were done. Higher IL-10 concentration was associated with increased egg counts (R^2^=0.1948: p=0.0146) **Figure 8D**.

**Figure 8.**
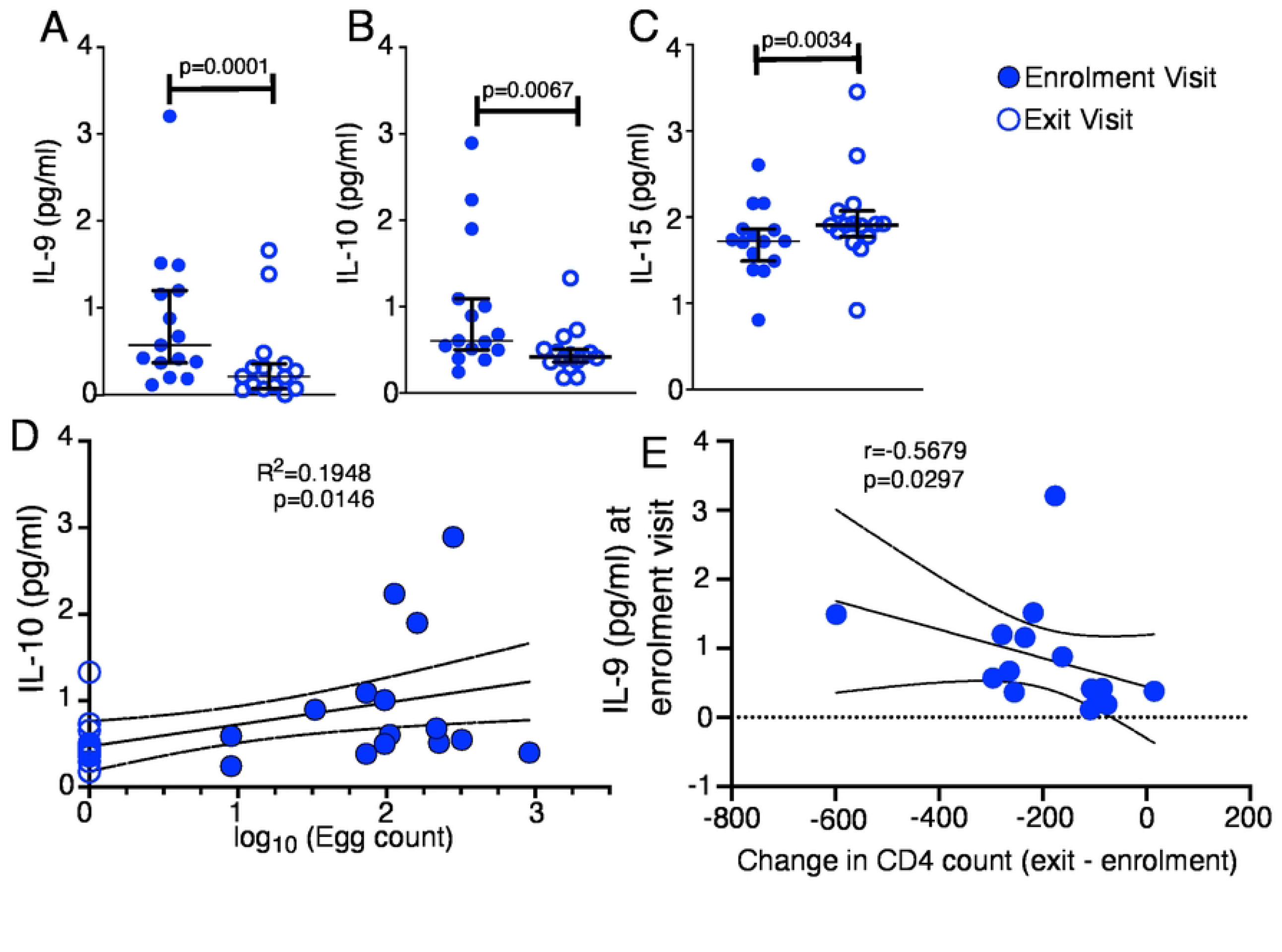
Effect of *S. mansoni* infection on IL-9 (A), IL-10 (B) and 1L-15 (C) plasma concentration. Each dot represents a cytokine concentration from a participant at either enrolment visit (filled) or exit visit (empty). The horizontal lines represent the mean and 95% confidence interval (C.I). Wilcoxon matched pairs signed rank test (A, B, C) and linear regression (D) and correlation (E) tests were used for the comparisons. n=17.

### IL-9 concentration at enrolment visit predicts CD4 decline

The difference in CD4 count (CD4 count at exit visit minus CD4 count at enrolment visit) was determined and associated with IL-9 concentration at enrolment visit. IL-9 concentration at enrolment visit negatively correlated with the difference in CD4 count between exit and enrolment visit (r=-0.5679 95% CI=-0.8417 to -0.06176, p=0.0297) **Figure 8E**.

## Discussion

The study was conducted between August 2012 and September 2015. During this time, the government policy was that only individuals with CD4 count below 350 CD4 cells per μl of blood were eligible for ART. However, this policy has since changed and any individual who tests HIV positive is eligible for ART irrespective of CD4 count.

In HIV and *S. mansoni* mixed infection, each pathogen is characterized by different cytokine secretion profiles, it is not clear whether the interaction is more detrimental or beneficial for the host when compared with each infection on its own.

Although the MCV and MCH were significantly higher at the exit visit compared to the enrolment visit, these were within the normal ranges.

Therefore, to understand the effect of *S. mansoni* infection on HIV specific responses, we compared HIV specific and nonspecific immune responses within an individual when HIV and *S. mansoni* coinfected and after *S. mansoni* was cleared with praziquantel treatment. We found that the frequency of HIV specific T cells within three cell clusters were higher when the participants were HIV only infected compared to when the participants were HIV and *S. mansoni* coinfected. The clusters had high expression intensities of IFN-γ and intermediate expression intensities of TNF-α, CD8 and CD3 and low expression intensities of perforin, TLR7, CD4, and IL-2 (cluster 11), intermediate expression intensities of CD8 and perforin and low expression intensities of IFN-γ, TNF-α, CD3, TLR-7, CD4 and IL-2 (cluster 16) and intermediate expression intensities of CD8 and perforin and low expression intensities of IFN-γ, TNF-α, CD3, TLR7, CD4 and IL-2 (cluster 29). No other cluster of cells had significant higher or lower frequency of cells at either enrolment visit or exit visits. In addition, IL-9 and IL-10 concentration were lower and IL-15 was higher when the participants were HIV only infected compared to when the participants were HIV and *S. mansoni* coinfected. The intermediate and low expression of CD3 in all these clusters is likely due to downregulated CD3 upon stimulation in responders.

The cells in clusters 11, 16 and 29 were predominantly CD8 T cells. Cells in cluster 11 predominantly secreted IFN-γ, cells in cluster 16 and 29 predominantly secreted perforin. The PTE peptides are short and likely did not favor CD4 T cell stimulation. Together, the data suggests that *S. mansoni* infection enhanced downmodulation of HIV specific responses which recovered after *S. mansoni* clearance.

Immunoregulation function of IL-10 has been proposed as a mechanism through which *S. mansoni* induces downmodulation of Th1 immune responses [20, 21]. This study obtained similar results, where the IL-10 concentration is higher and IL-15 concentration is lower in the presence of *S. mansoni* infection. However, upon the clearance of *S. mansoni* infection by praziquantel treatment, IL-10 concentration reduced and IL-15 concentration increased. Furthermore, the *S. mansoni* egg burden increased with increase in IL-10 production.

IL-10 has been shown to limit antigen specific responses, as a means to balance between immune responses and pathology, however, in the presence of *S. mansoni* infection, higher concentrations of IL-10 are secreted, suggesting an enhanced immunoregulation to non S. *mansoni* immune responses including HIV.

In our study, *S. mansoni* infection impaired IL-15 secretion in the plasma of HIV and *S. mansoni* coinfected individuals. IL-15 has been shown to stimulate anti-HIV immunity by enhancing CD8 T cell and NK cell functionality [22, 23]. IL-15 response signature predicts Rh CMV/SIV vaccine efficacy [24].

IL-9 plays a significant role in expelling worms [25] and in the absence of the worms at the exit visit, IL-9 responses declined to basal level. Interestingly, IL-9 concentration at enrolment predicted greater CD4 T cell decline in HIV and *S. mansoni* coinfected individuals suggesting an enhanced CD4 loss. The data suggests that IL-9 may be playing a role with the delayed replenishment of CD4 cells after cell death.

Importantly, our study establishes an immune responses mechanism in a mixed infection of HIV and *S. mansoni*, where the interactions of these two pathogens downmodulates the immune responses involved in the control HIV. These data highlight the impairment of HIV specific immunity in co-infections and stress the need to develop combination therapies.

Due to study design constraints, we were not able to identify the cells that secreted IL-9 in the plasma.

## Abbreviations

PTE: Potential T cell epitope
PBS-T: PBS plus 0.05% Tween-20
PZQ: Praziquantel
ART: Antiretroviral therapy
UMAP: Uniform Manifold Approximation and Projection.
pVL: Plasma viral load

## Acknowledgement

The authors acknowledge Ben Gombe for PBMC isolation and sample shipment and William Senyonga for technical assistance in the CAA assays. We also acknowledge Pietro Pala for giving insights into study planning.

